# Long non-coding RNA Gm15441 attenuates hepatic inflammasome activation in response to metabolic stress

**DOI:** 10.1101/675785

**Authors:** Chad N. Brocker, Donghwan Kim, Tisha Melia, Kritika Karri, Thomas J. Velenosi, Shogo Takahashi, Jessica A. Bonzo, David J. Waxman, Frank J. Gonzalez

**Author notes:** These authors contributed equally (Chad N. Brocker, Donghwan Kim).

## Abstract

Fasting paradigms elicit a wide-range of health benefits including suppressing inflammation. Exploring the molecular mechanisms that prevent inflammation during caloric restriction may yield promising new therapeutic targets. During fasting, activation of the nuclear receptor peroxisome proliferator-activated receptor alpha (PPARA) promotes the utilization of lipids as an energy source. Herein, we show that ligand activation of PPARA directly upregulates the long non-coding RNA gene *Gm15441* through binding sites within its promoter. *Gm15441* expression suppresses its antisense transcript, encoding thioredoxin interacting protein (TXNIP). This, in turn, decreases TXNIP-stimulated NLRP3 inflammasome activation, caspase-1 (CASP1) cleavage, and proinflammatory interleukin 1 beta (IL1B) maturation. *Gm15441*-null mice were developed and shown to be more susceptible to NLRP3 inflammasome activation and to exhibit elevated CASP1 and IL1B cleavage in response to metabolic and inflammatory stimuli. These findings provide evidence for a novel mechanism by which PPARA attenuates hepatic inflammasome activation in response to metabolic stress through lncRNA *Gm15441* induction.

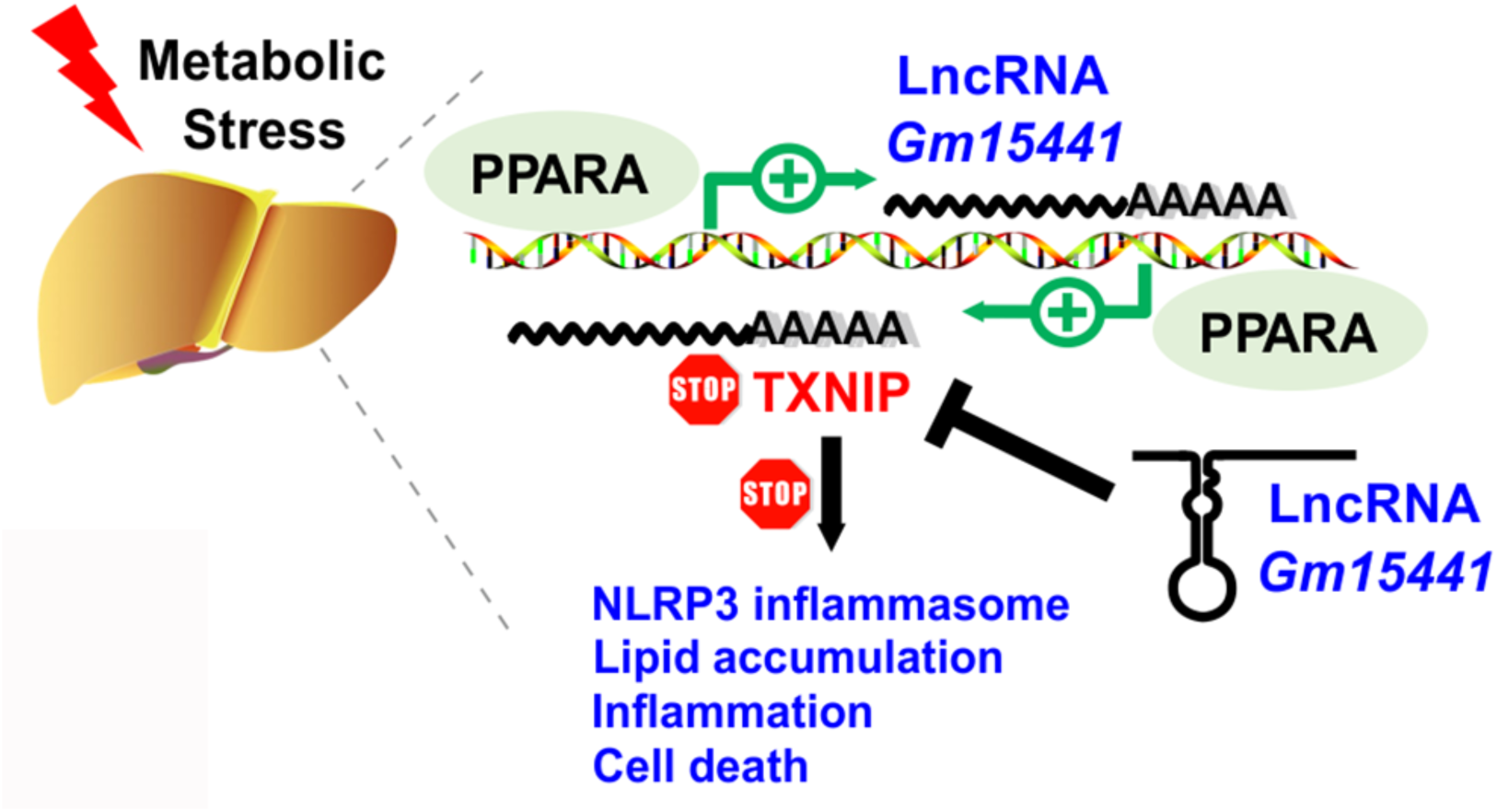
Graphical abstract

## Introduction

There is growing support for a strong link between metabolism and inflammation (Andrejeva and Rathmell, 2017; Bettencourt and Powell, 2017). Various fasting regimes are known to provide many health benefits including anti-inflammatory effects (Corrales et al., 2019). Peroxisome proliferator-activated receptor alpha (PPARA) is a ligand-activated nuclear receptor and transcription factor that is a key regulator of the fasting response. PPARA facilitates metabolic remodeling that promotes lipid oxidation, and its dysregulation contributes to metabolic disorders and liver disease (Brocker et al., 2018). Further, synthetic PPARA agonists act as potent anti-inflammatory agents (Yang et al., 2017; Zheng et al., 2017). However, the regulatory mechanisms underlying how PPARA prevents inflammation are not well understood.

Long non-coding RNAs (lncRNAs) act as regulators of gene expression and play important regulatory roles in many metabolic processes. For example, lncRNA Blnc1 works in concert with another important hepatic nuclear receptor, liver X receptor (LXR), to activate the lipogenic gene program (Zhao et al., 2018). Down-regulation of lncRNA lncOb reduces leptin, leading to a leptin responsive form of obesity (Dallner et al., 2019). Other studies found that lncRNAs are regulated during adipocyte differentiation (Yuan et al., 2019a; Yuan et al., 2019b), are expressed in liver in a sex-dependent manner when regulated by growth hormone (Melia et al., 2016; Melia and Waxman, 2019), and can be strongly induced by xenobiotic exposure (Lodato et al., 2017). A compartment-specific transcriptional profiling approach revealed that lncRNA PAXIP1-AS1 regulates pulmonary arterial hypertension by modulating smooth muscle cell function (Jandl et al., 2019). Further, lncRNA HOTAIR influences glucose metabolism by upregulation of GLUT1 in hepatocellular carcinoma cells (Wei et al., 2017). It is therefore reasonable to consider that lncRNAs may play important roles in the metabolic remodeling and anti-inflammatory actions that occur after PPARA activation.

In the present study, the mechanisms by which PPARA prevents inflammation during periods of metabolic stress were investigated. RNA-seq was carried out on livers from mice treated with the PPARA agonist WY-14643, and large numbers of differentially expressed protein coding genes and lncRNA genes were identified. These studies led to discovery of a regulatory axis between thioredoxin-interacting protein (TXNIP) and an antisense lncRNA, *Gm15441*. TXNIP acts as a critical relay linking oxidative and endoplasmic reticulum (ER) stress to inflammation through NLRP3 inflammasome activation (Anthony and Wek, 2012; Oslowski et al., 2012). Studies have also shown that NLRP3 inflammasome activity is attenuated by mono- and polyunsaturated fatty acids, which are endogenous PPARA agonists (Forman et al., 1997; Kliewer et al., 1997; Mochizuki et al., 2006; Ralston et al., 2017; Shen et al., 2017). Considering these studies, a regulatory mechanism was proposed whereby fatty acids mobilized from adipose tissue during fasting activate PPARA, which in turn suppresses the NLRP3 inflammasome by strong induction of the TXNIP-suppressing lncRNA gene *Gm15441*, which is antisense to *Txnip*. ChIP-seq datasets were analyzed for PPARA binding sites within the *Gm15441* promoter and *Gm15441* regulatory elements were confirmed using reporter gene assays and PPARA ChIP studies. *Gm15441* transgene expression downregulated *Txnip*, demonstrating lncRNA-mediated gene suppression *in vitro*. CRIPSR/Cas9-mediated gene editing was employed to develop a *Gm15441* knockout (*Gm15441*^LSL^) mouse model. Basal TXNIP levels in *Gm15441*^LSL^ mice were significantly elevated over wild-type mice levels, supporting *Gm15441* as a negative regulator of *Txnip* expression *in vivo*. *Gm15441*^LSL^ mice were treated with a PPARA agonist or fasted to assess how loss of *Gm15441* impacts hepatic inflammasome activation in response to both pharmacological and physiologically-induced metabolic stress. TXNIP protein, caspase-1 (CASP1) levels, and interleukin 1 beta (IL1B) cleavage were elevated in *Gm15441*^LSL^ mice and were further increased by PPARA activation, indicating that this lncRNA plays a major role in attenuating inflammasome activation. Thus, hepatic PPARA directly regulates the lncRNA *Gm15441*, which in turn suppresses *Txnip* expression, which attenuates NLRP3 inflammasome activation during periods of metabolic stress. These studies revealed a novel regulatory mechanism supporting the beneficial effects of fasting, namely, reduced inflammation.

## Results

### LncRNA regulation by PPARA is highly tissue-specific

To assess the regulation of lncRNAs by PPARA, RNA-seq was performed using total liver RNA isolated from *Ppara*^+/+^ mice, both with and without dietary exposure to the PPARA agonist WY-14643. A lncRNA discovery pipeline was implemented that identified 15,558 liver-expressed lncRNA genes. Of these, 13,343 were intergenic, 1,966 were antisense, and 249 were intragenic lncRNAs. 44% of the 15,558 liver-expressed lncRNAs are considered novel (Melia and Waxman, 2019). Differential gene expression analysis revealed that a total of 1,735 RefSeq genes and 442 liver-expressed lncRNA genes responded to treatment with WY-14643 at an expression fold-change >2 at FDR<0.05, with 968 RefSeq genes and 245 lncRNA genes upregulated, and 767 RefSeq genes and 197 lncRNA genes downregulated following WY-14643 treatment (**Table S1A and Table S1B**). Only 17 RefSeq genes (1.0%) and 6 lncRNAs (1.3%) responded to WY-14643 in the same manner in PPARA-knockout mice. Thus, ∼99% of these transcriptomic responses are PPARA-dependent (**Figure 1A**).

**Figure 1.**
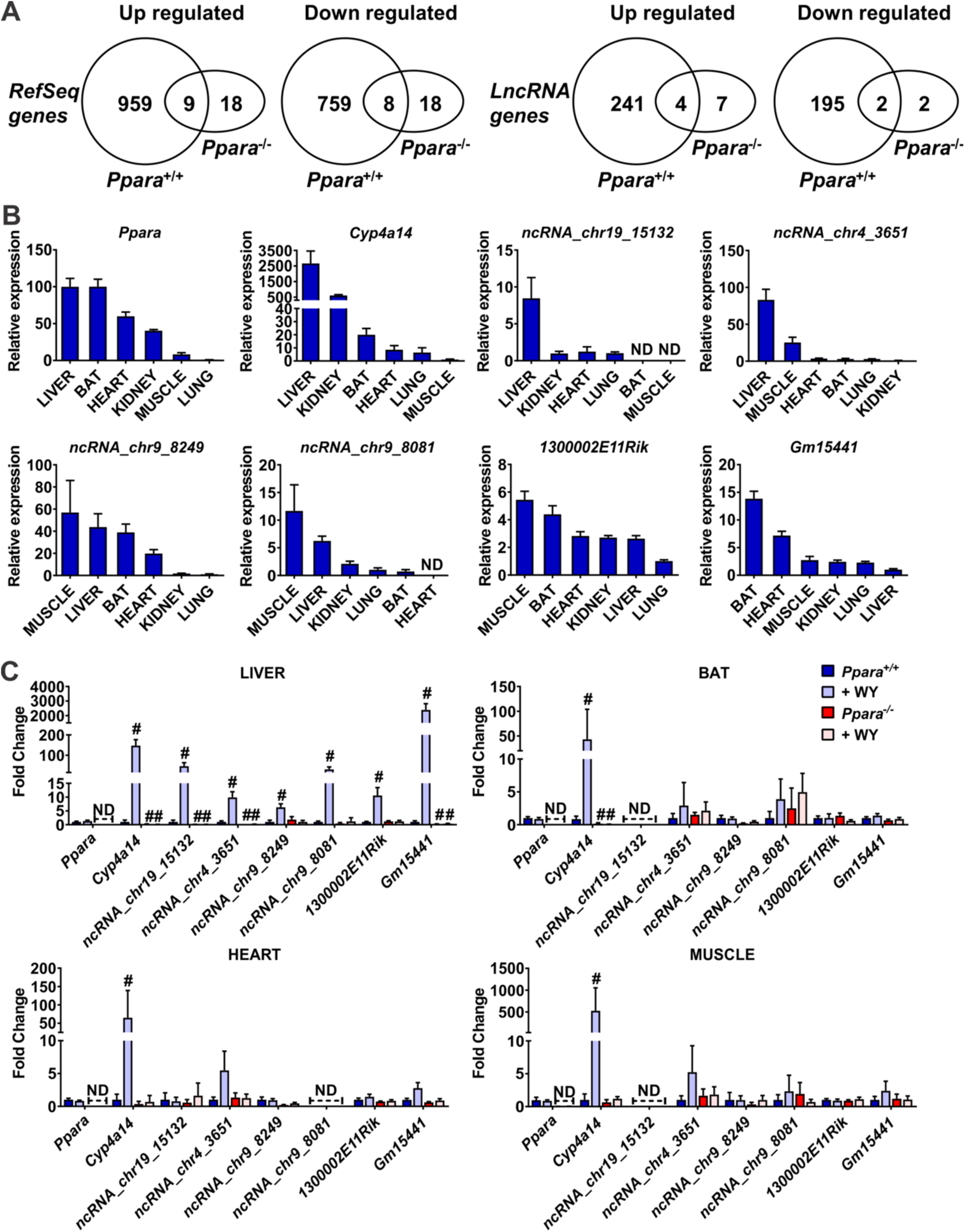
LncRNA identified as PPARA targets are expressed in several oxidative tissues but exhibit liver-specific transcriptional response to WY-14643. (A) Venn diagrams of RefSeq transcripts and lncRNA transcripts that were differentially regulated by WY-14643 (fold change >2 at FDR < 0.05) in wild-type and in *Ppara*^−/−^ mouse liver, as determined by RNA-seq. 123 of the RefSeq genes are non-coding, indicated by their NR accession numbers. Six RefSeq genes and six lncRNA genes show opposite responses to WY-14643 treatment and are excluded from the gene counts shown. (B) Relative basal lncRNA expression in select tissues. (C) LncRNA expression in tissues from *Ppara*^+/+^ and *Ppara*^−/−^ mice treated with WY-14643 for 48 hours. At least five mice were analyzed for each genotype and treatment group. Each data point represents the mean ± SD for n = 5 tissue samples. # *P* < 0.05. Abbreviation: ND, Not Detected.

Comparison of the PPARA-responsive lncRNAs to lncRNAs responsive to agonist ligands of two other nuclear receptors in mouse liver, namely CAR and PXR (Lodato et al., 2017), identified 8 lncRNAs that are induced by all three nuclear receptors and 11 lncRNAs repressed by all three receptors. Forty five other lncRNAs were induced, or were repressed, in common by PPARA and CAR, and 30 other lncRNAs were either induced or repressed in common by PPARA and PXR (**Table S1C**). Overall, 94 (21%) of the 442 PPARA-responsive liver-expressed lncRNAs showed common responses to CAR and/or PXR activation. Furthermore, 59 other PPARA-responsive lncRNAs showed opposite patterns of response to PPARA as compared to CAR or PXR (**Table S1C**). These findings are consistent with the partial functional overlap between these three xenobiotic nuclear receptors and their gene targets (Cui and Klaassen, 2016; Gonzalez et al., 2015; Wada et al., 2009).

Pathway analysis revealed that the WY-14643 upregulated genes are most highly enriched for the following biological processes: lipid metabolism, peroxisome, DNA replication, DNA repair, cell cycle and fatty acid oxidation (**Table S1D**). The downregulated genes are most highly enriched for: secreted factors, monooxygenase, immunity, serine protease inhibitor, metabolism of xenobiotics, and response to virus (**Table S1E**). PPARA-dependent lncRNAs identified by RNA-seq were selected and mRNA levels monitored over a 24 hour period after WY-14643 treatment, revealing that lncRNA expression profiles paralleled those of known protein-coding PPARA target genes (**Figure S1A-D**). As lncRNA expression is often highly tissue specific (Iwakiri et al., 2017; Melia et al., 2016; Perron et al., 2017), the relative basal levels were measured for PPARA-responsive lncRNAs expressed in oxidative tissues that utilize fatty acids as an energy source and express PPARA at the highest levels. *Ppara* mRNA and its classic target gene *Cyp4a14* served as controls. Basal levels of *Cyp4a14* mRNA and intergenic lncRNAs *ncRNA_chr19_15132* and *ncRNA_chr4_3651* were highest in liver, while *ncRNA_chr9_8249, ncRNA_chr9_8081*, and *1300002E11Rik* were most highly expressed in muscle. One of the PPARA-dependent lncRNAs, *Gm15441*, was also most highly expressed in brown adipose tissue (BAT) (**Figure 1B**).

To assess the impact of PPARA activation on lncRNA expression and tissue specificity, *Ppara* wild-type mice (*Ppara*^+/+^) and *Ppara*-null mice (*Ppara*^−/−^) were treated with WY-14643 for 48 hours, and four tissues (liver, BAT, heart, muscle) then harvested for analysis. *Cyp4a14* mRNA was markedly induced by WY-14643 in all four tissues and in a *Ppara*-dependent manner, as expected (**Figure 1C**). All six lncRNAs were induced in liver, including *Gm15441*, which exhibited >2000-fold increase in wild-type livers, > 10-fold higher than the other lncRNAs. In contrast to the induction of *Cyp4a14* mRNA, lncRNA induction by WY-14643 was liver-specific, with no significant induction of the five other lncRNAs seen in BAT, heart, or muscle (**Figure 1C**). Further, none of these lncRNAs was induced in *Ppara*^−/−^ mice in any tissue examined. Thus, PPARA-mediated induction of these lncRNAs is highly liver-specific, with *Gm15441* showing the most robust induction response.

### LncRNA *Gm15441* is antisense to *Txnip* and exhibits inverse regulation in response to PPARA activation

LncRNA *Gm15441* overlaps the coding region of *Txnip*, which is located on the opposing strand and also responds to WY-14643 treatment, albeit with different kinetics than *Gm15441*. Analysis of the mapped sequence reads obtained by stranded RNA sequencing revealed a pronounced increase in expression of *Gm15441* (**Figure 2A**), whereas, the *Txnip* transcript on the opposite strand was suppressed by WY-14643 at the time point analyzed (**Figure 2B**). qRT-PCR analysis confirmed the very large, highly significant increase in *Gm15441* expression and its complete inhibition in *Ppara^−/−^* liver (**Figure 2C**). Further, *Txnip* RNA was significantly suppressed by WY-14643 in wild-type mice (**Figure 2D**). Additionally, *Txnip* expression was attenuated in untreated *Ppara^−/−^* compared to *Ppara^+/+^* mouse liver, indicating a role for PPARA in maintaining basal expression of *Txnip* (**Figure 2D**). Given that *Txnip* expression can be upregulated in human neuroblastoma cells by the PPARA activator fenofibrate, which suppresses proliferation and migration (Su et al., 2015), the effects of a single dose of WY-14643 on *Gm15441* and *Txnip* mRNA were examined over a 24 hour period. *Txnip* mRNA was induced rapidly, with maximum expression seen at 1.5 hours, followed by a marked decrease coincident with the increased expression of *Gm15441* (**Figure 2E**). This time course is markedly different from that of 14 other WY-14643-inducible protein coding and lncRNA genes examined, where peak induction occurred 6 to 12 hours after WY-14643 treatment (**Figure S1**), as was also found for *Gm15441*. The unusual kinetics seen with *Txnip* – early induction followed by repression coinciding with the activation of *Gm15441 –* indicate that *Txnip* is a PPARA target gene whose expression in liver is inversely regulated with that of *Gm15441* after PPARA activation.

**Figure 2.**
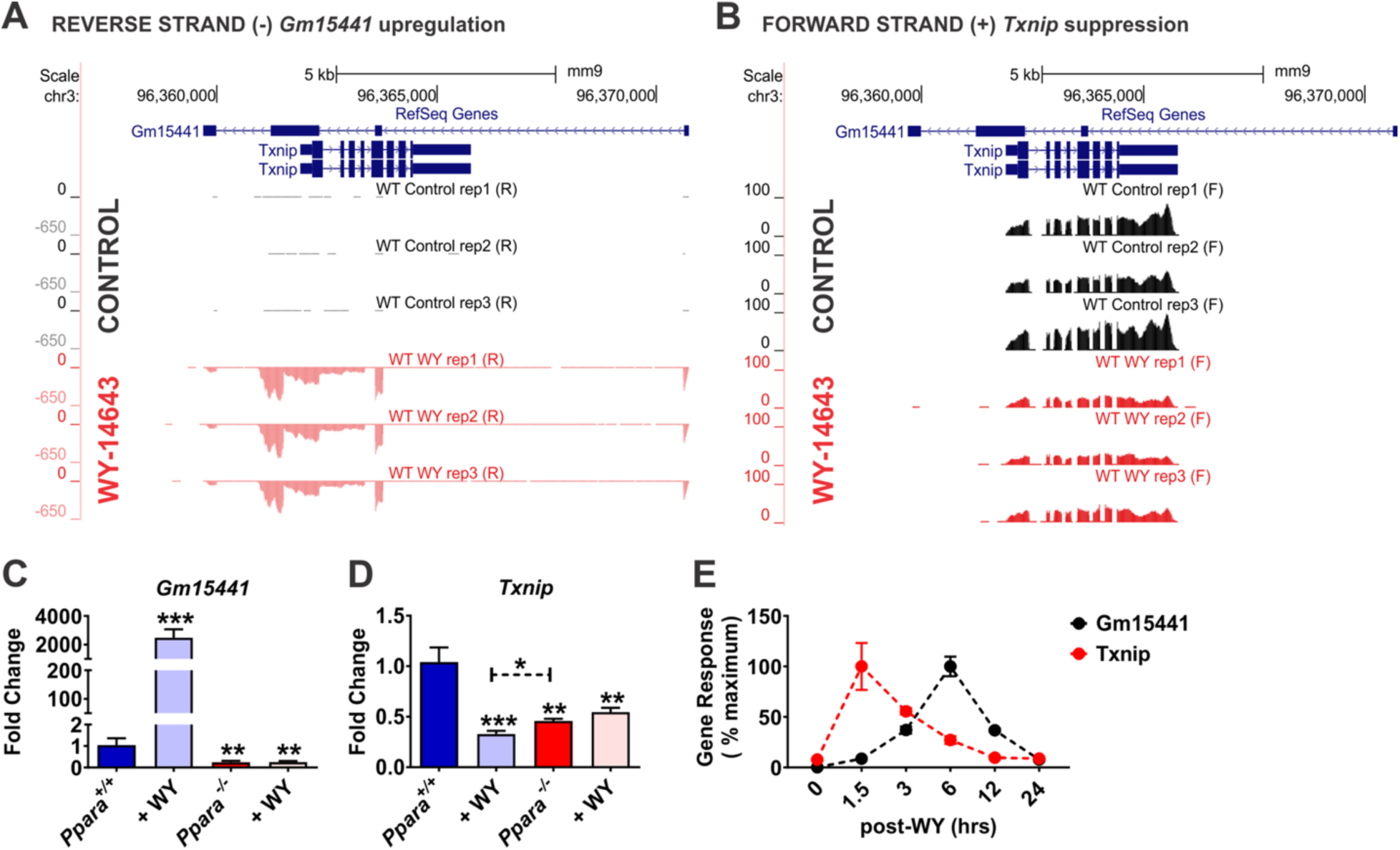
*Txnip* and lncRNA *Gm15441* are inversely regulated following PPARA activation. Wild-type (*Ppara*^+/+^) mice were treated with WY-14643 and stranded RNA-seq was performed on total liver mRNA. (A) Expression of the antisense lncRNA *Gm15441* is strongly upregulated. (B) Expression of the protein-coding *Txnip* mRNA is downregulated. Shown are the changes in expression of *Gm15441* (C) and *Txnip* (D) mRNAs in *Ppara*^+/+^ and *Ppara*^−/−^ mice treated with WY-14643, or vehicle control, for 48 hours, determined by qRT-PCR. (E) Time course for changes in expression of *Gm15441* and *Txnip* mRNA over a 24 hours period following treatment with WY-14643 by gavage, determined by qRT-PCR. The maximum response of *Txnip* mRNA was seen at 1.5 hours and for *Gm15441* was seen at 6 hours. Each data point represents the mean ± SD for n = 5 liver samples. *P < 0.05; **P < 0.01; ***P < 0.001.

### PPARA directly regulates lncRNA *Gm15441* by binding to its promotor

To identify PPARA binding sites at the *Gm15441*/*Txnip* locus, PPARA ChIP-seq datasets (GSE61817) from PPARA agonist-treated mice (Lee et al., 2014a) were analyzed (**Figure 3A**). Three major ChIP-seq peaks indicating PPARA binding were seen upstream of *Gm15441*, and one major peak was seen in the promoter region of *Txnip.* Examination of genomic sequences upstream of *Gm15441* using Genomatix MatInspector (Genomatix, Munchen, Germany) identified seven peroxisome proliferator response elements (PPREs) within six genomic regions, designated A to F, within 10 kilobase (kb) upstream of the *Gm15441* transcriptional start site (TSS) (**Figure 3B**). These six *Gm15441* upstream sequences were synthesized and cloned into the pGL4.11 reporter, and luciferase assays were performed to assay the functionality of PPARA binding to the *Gm15441* promoter. A PPRE-luciferase construct containing an *Acox1* PPRE repeat was used as a positive control, and an empty pGL4.11 plasmid was used as a negative control. Luciferase activity was significantly elevated in primary mouse hepatocytes transfected with five of the six pGL4.11 constructs, consistent with direct regulation by PPARA at multiple loci (**Figure 3C**). PPARA binding was also assessed by ChIP assays using a polyclonal PPARA antibody and chromatin isolated from livers of wild-type and *Ppara*^−/−^ mice fed either control diet or a diet-containing WY-14643. Enrichment of PPARA binding to PPREs of known PPARA target genes, namely *Acot1* and *Acoxt1*, was determined by comparing binding to liver chromatin from wild-type mice fed control diet vs. WY-14643-containing diet (**Figure 3D**). *Ppara*^−/−^ livers were used as a negative control to identify non-specific binding. *Fscn2* primers were used as a non-target gene promoter and negative control. Enrichment of PPARA binding was found using primer sets covering *Gm15441* regions B, C, D, and F, while no binding was seen with *Gm15441* region E (**Figure 3D**), which was transcriptionally inactive (**Figure 3C**). Enrichment was strongest with region C, in agreement with the luciferase reporter gene data. Enrichment at sites C and D was significantly increased by WY-14643 treatment, indicating increased PPARA recruitment at these sites. No PPARA binding was seen with chromatin from *Ppara*^−/−^ mice on the control diet. Together, these data indicate that PPARA regulates *Gm15441* transcription by direct binding of PPARA to multiple PPRE sites within 10 kb of the *Gm15441* TSS.

**Figure 3.**
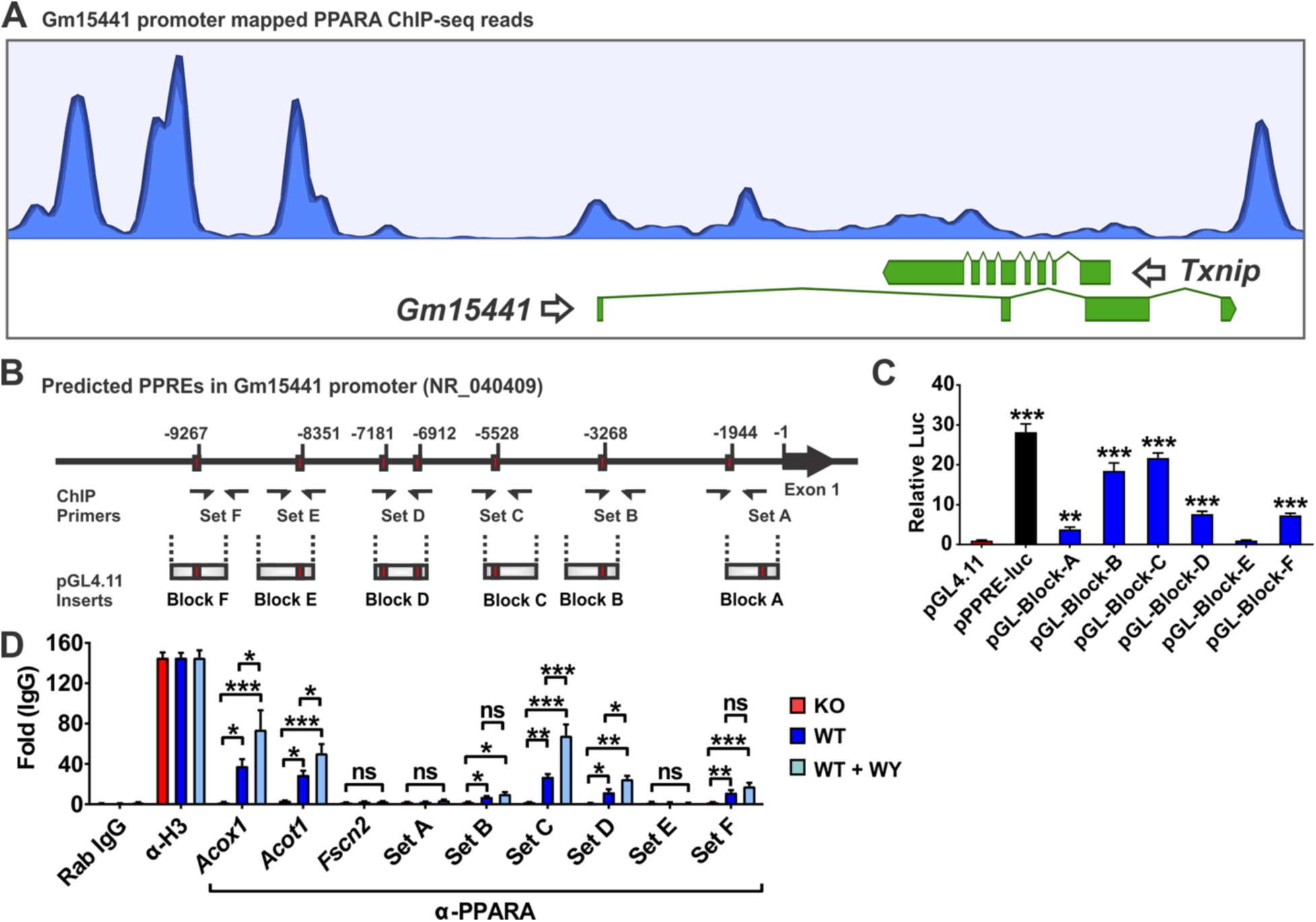
LncRNA *Gm15441* is a direct PPARA target gene. (A) PPARA ChIP-seq read peaks from agonist (GW7647)-treated mouse liver. (B) Schematic representation of seven PPRE sequences found within the *Gm15441* promoter (−10 kb). ChIP primer binding sites and reporter gene construct inserts are shown. (C) Luciferase-based reporter assays identified five functional PPREs within the *Gm15441* promoter, based on n = 3 replicates. (D) PPARA ChIP assays assessed PPRE binding in liver samples from *Ppara*^+/+^ and *Ppara*^−/−^ mice treated with WY-14643. Experiments were performed with at least four different livers. Rabbit IgG and antibody to histone protein H3 were used as negative and positive controls, respectively. Each data point represents the mean ± SD for n = 5 liver samples. *P < 0.05; **P < 0.01; ***P < 0.001 for comparisons to pGL4.11 empty vector (C) or as indicated (D). Abbreviation: ns, not significant.

### Generation of a strand-specific Gm15441 knockout mouse

A gene targeting strategy was developed to generate a *Gm15441* knockout mice without impacting *Txnip* expression on the opposing strand. To accomplish this goal, CRISPR/Cas9 was used to insert a lox-STOP-lox (LSL) cassette downstream of *Gm15441* exon 1 (**Figure 4A and B**). The LSL cassette prevents transcription of *Gm15441* in a *Gm15441*-floxed mouse but has no effect on transcription of *Txnip*, which is > 2 kb downstream of the last exon of *Txnip*. Crossing with a transgenic *Cre* mouse deletes the LSL cassette and restores *Gm15441* expression (**Figure 4C**). All mouse lines were on a pure C57BL/6J background. The genotyping scheme generates an approximately 1200 bp band in *Gm15441* wild-type mice (*Gm15441*^+/+^) and a 1700 bp band in *Gm15441*^LSL^ mice (**Figure 4D**). Basal expression of *Gm15441* was significantly lower in livers of *Gm15441*^LSL^ when compared to *Gm15441*^+/+^ mice (**Figure 4E**). Expression in heterozygous *Gm15441*^HET^ mice was comparable to *Gm15441*^LSL^. To verify the subcellular localization and loss of expression of *Gm15441*, fluorescence *in situ* hybridization (FISH) staining was performed on livers from *Gm15441*^+/+^ and *Gm15441*^LSL^ mice fed control diet or treated with WY-14643. DAPI staining was used as a counter stain to detect nuclei in liver sections. A pronounced increase in Gm15441 fluorescence was detected in the cytoplasm and nuclei of hepatocytes in livers from WY-14643-treated *Gm15441*^+/+^ mice. The Gm15441 RNA signal was absent in livers from vehicle-treated *Gm15441*^+/+^ and *Gm15441*^LSL^ mice, and in livers from WY-14643-treated *Gm15441*^LSL^ mice (**Figure 4F**). These data validate *Gm15441*^LSL^ mice as an effective knockout mouse model and reveal that hepatic *Gm15441* expression is both nuclear and cytosolic evoking several possible mechanisms by which it regulates *Txnip*.

**Figure 4.**
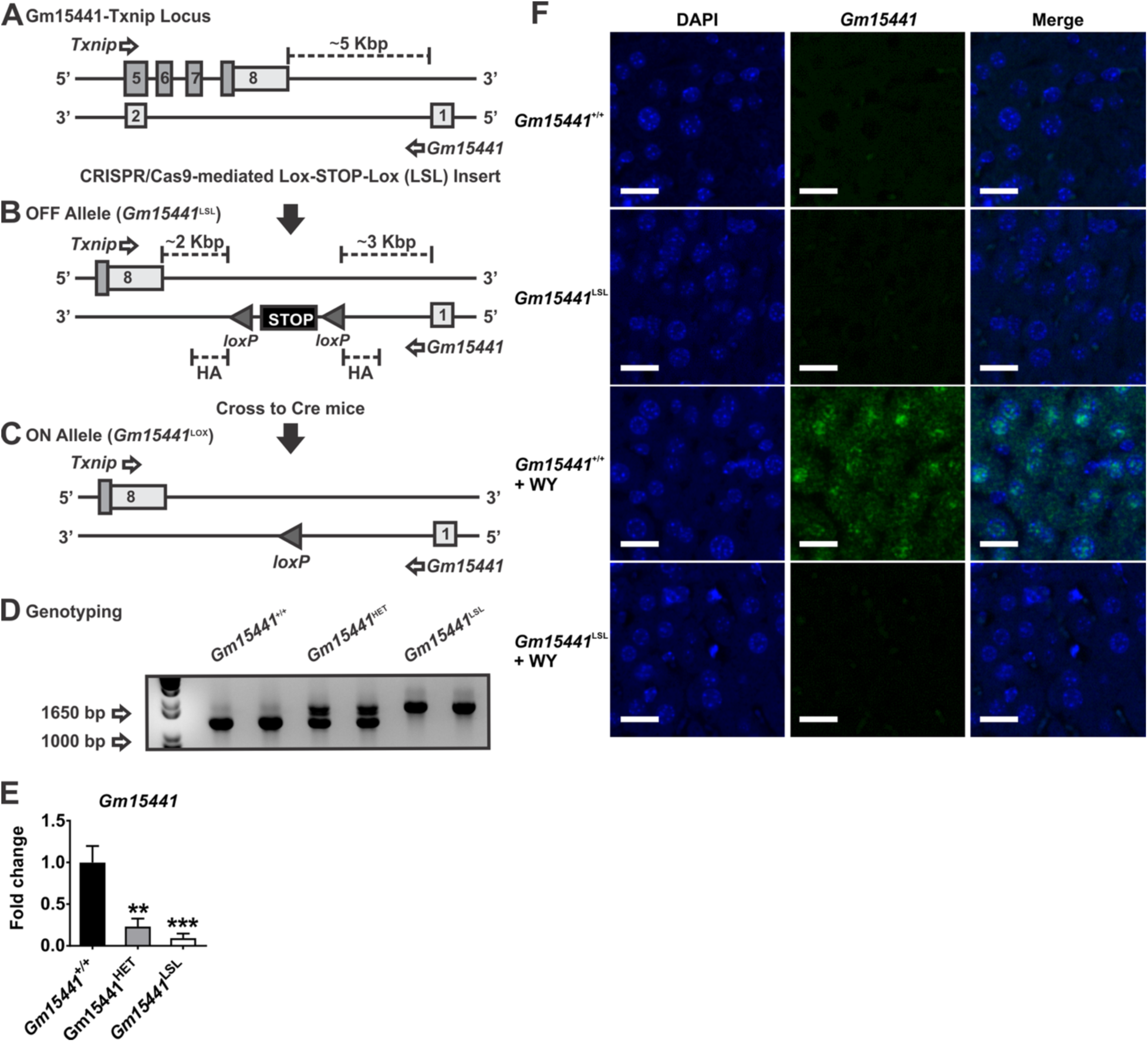
Generation of the *Gm15441*-null mouse line. Targeting strategy for generating a strand-specific *Gm15441*-null mouse line. (A) Exon structure of targeted *Gm15441*-*Txnip* locus. (**B**) CRISPR/Cas9-mediated insertion of Lox-STOP-Lox (LSL) cassette selectively ablates *Gm15441* expression (**C**) Cre-mediated removal of STOP cassette rescues *Gm15441* expression. (**D**) *Gm15441* knockout mouse genotyping. (**E**) Analysis of *Gm15441* mRNA by qRT-PCR from livers of *Gm15441*^+/+^, *Gm15441*^+/−^, and *Gm15441*^LSL^ mice. Each data point represents the mean ± SD for n = 5 liver samples. **P < 0.01; ***P < 0.001. (**F**) Fluorescence in situ hybridization staining of Gm15441 RNA in livers of *Gm15441*^+/+^ and *Gm15441*^LSL^ mice treated with WY-14643 for 48 hours. Scale bars represents 20 nm (200x).

### Loss of *Gm15441* potentiates inflammasome activation by WY-14643-induced metabolic stimulation

PPARA plays a role in attenuating inflammation in many tissues and disease models (Kono et al., 2009; Schaefer et al., 2008; Yoo et al., 2013; Yoo et al., 2011). In the liver, PPARA ameliorates inflammation by reducing ER stress in hepatocytes (Zhang et al., 2016). TXNIP is known to facilitate ER stress-induced NLRP3 inflammasome activation (Anthony and Wek, 2012; Lee et al., 2014b; Lerner et al., 2012; Oslowski et al., 2012). By regulating TXNIP levels, Gm15441 could serve as a link between PPARA and the NLRP3 inflammasome. To ascertain whether lncRNA Gm15441 attenuates inflammasome activation by regulating TXNIP levels, mice were treated with WY-14643 for 48 hours, and weight loss, liver index, ALT, AST, and serum glucose levels were measured. There was no discernable difference in weight loss between *Gm15441*^+/+^ and *Gm15441*^LSL^ mice on a WY-14643 diet (**Figure 5A**). Serum ALT and AST levels were unchanged by genotype or following WY-14643 treatment. Serum glucose levels were significantly decreased by WY-14643 in both *Gm15441*^+/+^ and *Gm15441*^LSL^ mice, but the decrease was significantly greater in *Gm15441*^LSL^ mice (**Figure 5A**). Furthermore, upon gross liver examination, pronounced hepatomegaly was induced by WY-14643 treatment in both *Gm15441*^+/+^ and *Gm15441*^LSL^ mice (**Figure 5B**). Pronounced swelling of hepatocytes was noted in livers of WY-14643-treated *Gm15441*^+/+^ and *Gm15441*^LSL^ mice (**Figure 5B**). Expression of the PPARA target genes acyl-CoA dehydrogenase medium chain (*Acadm*) and *Cyp4a14* was significantly increased by WY-14643 in both *Gm15441*^+/+^ and *Gm15441*^LSL^ mice. *Gm15441* was strongly upregulated in livers of wild-type mice but was not detected in *Gm15441*^LSL^ mice. *Txnip* mRNA was significantly decreased by WY-14643 in livers of wild-type animals, consistent with Figure 2, but was significantly induced in *Gm15441*^LSL^ livers (**Figure 5C**). Basal expression of the inflammasome-related proteins TXNIP, cleaved caspase 1 (CASP1), and cleaved interleukin 1β (IL1B) was elevated in livers of *Gm15441*^LSL^ mice compared to wild-type mice, and significant further increases were seen following WY-14643 treatment (**Figure 5D, 5E**). Overall, levels of all three proteins were much higher in livers of WY-14643-treated *Gm15441*^LSL^ mice than wild-type mice (**Figure 5E**). Thus, ablation of *Gm15441* increases the expression of TXNIP and two other proteins associated with NLRP3 inflammasome activation, namely, cleaved CASP1 and IL1B.

**Figure 5.**
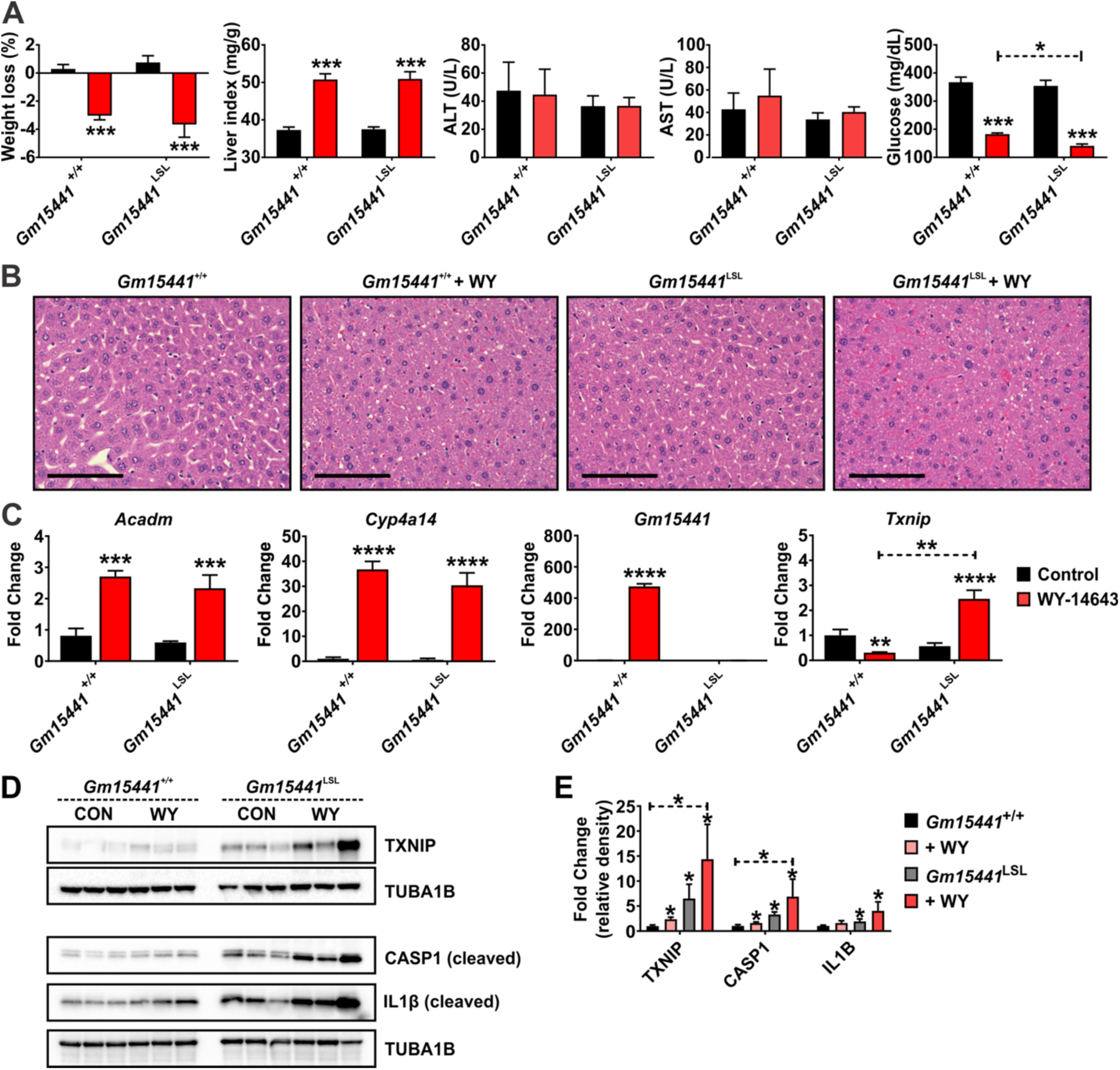
Loss of *Gm15441* potentiates inflammasome activation by WY-14643-induced metabolic stimulation. (A) Physiological endpoints from *Gm15441*^+/+^ and *Gm15441*^LSL^ mice treated with WY-14643 for 48 hours. Liver indexes (mg liver/g body mass) in response to WY-14643 treatment. (B) H&E staining of liver tissue. Scale bars represents 100 nm (400x). (C) Analysis of *Acadm*, *Cyp4a14*, *Txnip*, and *Gm15441* mRNAs in livers of *Gm15441*^+/+^ and *Gm15441*^LSL^ mice treated with WY-14643 for 48 hours, measured by qRT-PCR. (D) Analysis of TXNIP, TUBA1B, CASP1 (cleaved), and IL1B (cleaved) protein on livers from *Gm15441*^+/+^ and *Gm15441*^LSL^ mice treated with WY-14643 for 48 hours. (E) Densitometric analysis of TXNIP, CASP1, and IL1B protein levels. Each data point represents mean ± SD for n = 5 liver samples. *P < 0.05; **P < 0.01; ***P < 0.001; ****P < 0.0001 for comparisons between WY-14643-stimulated and unstimulated livers of the same genotype, or as shown (dashed horizontal lines).

### Loss of Gm15441 potentiates inflammasome activation during physiological response to acute fasting

During fasting, hepatic PPARA facilitates the metabolic remodeling that promotes use of lipids as an alternate energy source (Brocker et al., 2018). Fasting is also associated with many anti-inflammatory effects (Corrales et al., 2019). To determine whether *Gm15441* regulates inflammasome activation during physiological fasting, *Gm15441*^+/+^ and *Gm15441*^LSL^ mice were fasted for 24 hours. Serum ALT and AST were not changed between *Gm15441*^+/+^ and *Gm15441*^LSL^ mice, both with and without WY-14643 treatment (**Figure 6A**). Fasting decreased serum glucose levels in both *Gm15441*^+/+^ and *Gm15441*^LSL^ mice, but the decrease was greater in *Gm15441*^LSL^ mice (**Figure 6A**). Expression of *Acadm*, *Cyp4a14*, and *Txnip* mRNAs was significantly increased by fasting in both *Gm15441*^+/+^ and *Gm15441*^LSL^ mice (**Figure S2A**). *Gm15441* was significantly induced by fasting, although to a much lower degree than with WY-14643 treatment (**Figure S2A**). Expression of *Txnip* mRNA was increased to the same level in livers of fasted *Gm15441*^LSL^ mice as in fasted *Gm15441*^LSL^ mice (**Figure S2A**). Fasting also increased TXNIP, CASP1, and IL1B protein levels in *Gm15441*^LSL^ mice, in contrast to either a modest increase (TXNIP) or no increase (CASP1, IL1B) in *Gm15441*^+/+^ mice (**Figure S2B, Figure S2C**). Fasting markedly promotes lipid accumulation in mouse liver (Brocker et al., 2018). Lipid droplet accumulation was also observed and more widespread in fasting *Gm15441*^LSL^ mice than fasting *Gm15441*^+/+^ mice (**Figure S2D**). ORO staining was performed to validate the lipid accumulation response to fasting in *Gm15441*^LSL^ mice. A greater amount of lipid accumulation was observed in fasting *Gm15441*^LSL^ liver than in fasting *Gm15441*^+/+^ mice liver (**Figure 6B**). Serum and liver triglyceride (TG) levels and liver (but not serum) total cholesterol (CHOL) levels were significantly elevated in fasting *Gm15441*^LSL^ mouse livers compared to fasting *Gm15441*^+/+^ mouse livers (**Figure 6C**). These findings were supported by RNA-sequencing analysis, which revealed that liver damage and lipid accumulation-related genes were impacted by *Gm15441* deficiency, albeit in a complex manner (**Table S2**). Furthermore, analysis of the impact of fasting on mRNAs for apolipoprotein A4 (*Apoa4*), betaine-homocysteine methyltransferase (*Bhmt*), lipocalin 2 (*Lcn2*), orosomucoid 2 (*Orm2*), serum amyloid A1 (*Saa1*), and serum amyloid A2 (*Saa2*) – biomarker genes for liver damage, inflammation, and/or fatty accumulation – showed these genes were significantly elevated in fasting *Gm15441*^LSL^ compared to *Gm15441*^+/+^ mouse livers (**Figure 6D**). Fasting induced lower but still significant increases in several of these mRNAs in *Gm15441*^+/+^ livers, which could reflect the fasting-induced increase in *Txnip* expression (**Figure S2**). Thus, *Gm15441* negatively regulates the NLRP3 inflammasome pathway and lipid accumulation, potentially by preventing *Txnip* expression, in response to fasting.

**Figure 6.**
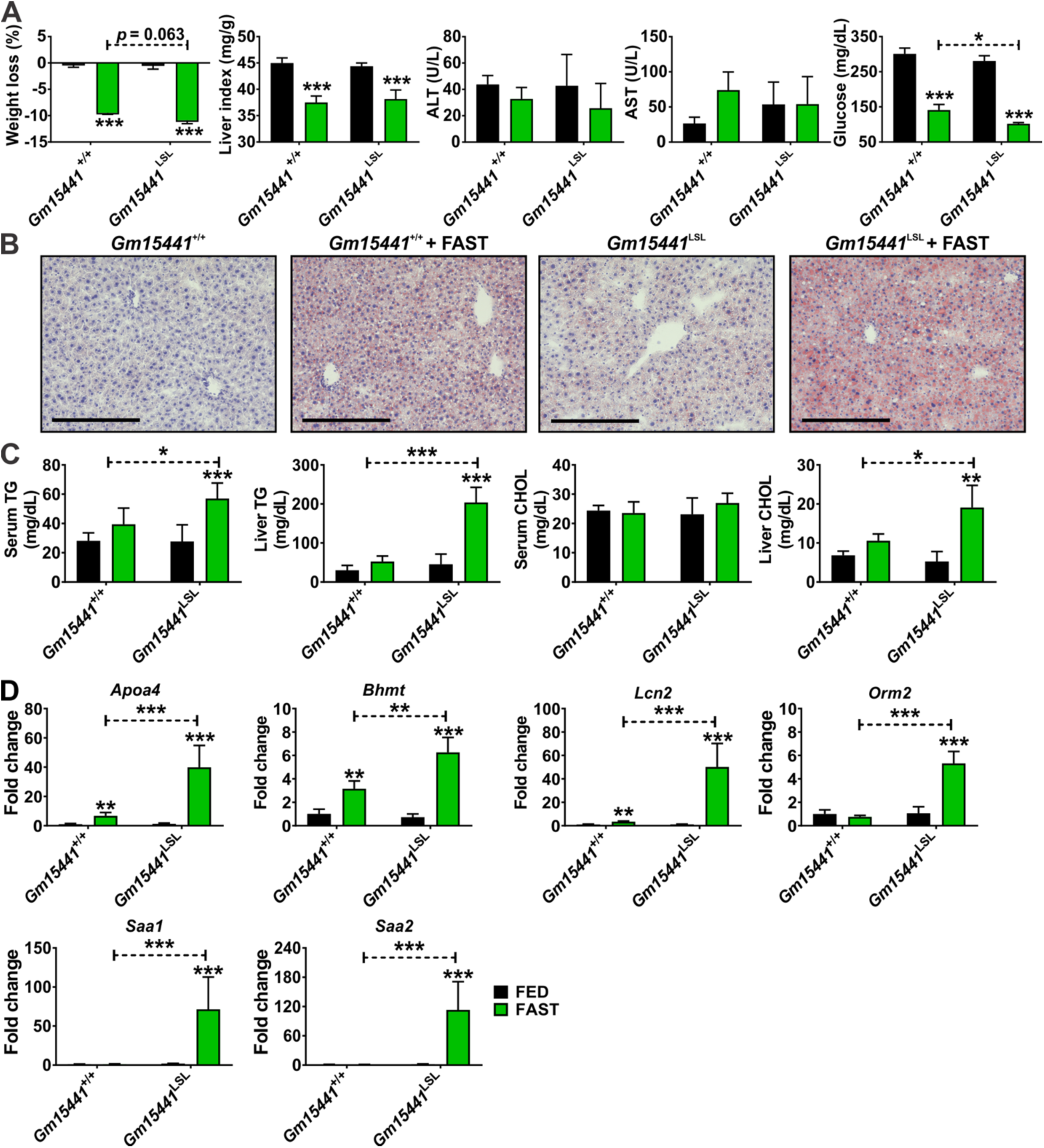
Loss of *Gm15441* potentiates inflammasome activation during physiological response to acute fasting. (A) Physiological endpoints from *Gm15441*^+/+^ and *Gm15441*^LSL^ mice after 24 hours fasting. Liver indexes (mg liver/g body mass) in response to fasting. (B) ORO staining of liver tissues after a 24 h fast. Scale bars represents 100 nm (200x). (C) TG and CHOL levels from serum and liver tissues after a 24 h fast. (D) Analysis of *Apoa4*, *Bhmt*, *Lcn2*, *Orm2*, *Saa1*, and *Saa2* mRNAs in livers of *Gm15441*^+/+^ and *Gm15441*^LSL^ mice after 24 h fast. Each data point represents the mean ± SD for n = 5 liver samples. *P < 0.05; **P < 0.01; ***P < 0.001 for comparisons between fast-stimulated and unstimulated livers of the same genotype, or as shown (dashed horizontal lines).

### Translational regulation of TXNIP by Gm15441 is partially dependent on 5’ UTR sequences

Translational regulation of TXNIP can occur via an internal ribosome entry site (IRES) sequences found in the 5’UTR (Lampe et al., 2018). IRES sequences are commonly associated with cell survival- and stress response-related genes and utilized when cap-dependent translation is inhibited during nutrient deprivation (Wu et al., 2014). The 5’UTR of *Txnip* contains multiple binding sites for polypyrimidine tract-binding protein (PTB), which regulates IRES-mediated translation (Lampe et al., 2018). As *Gm15441* transcription generates an RNA antisense to the 5’UTR of *Txnip*, *Gm15441* RNA may block PTB binding to the *Txnip* 5’UTR and thereby inhibit *Txnip* translation. To investigate this regulatory mechanism, the translational inhibitory potential of the *Txnip* 5’UTR was examined when linked to the coding sequence for GFP (**Figure 7A**). The non-5’UTR-GFP and the 5’UTR-GFP expression vectors were each transfected into Hepa-1 cells together with a *Gm15441* expression vector or an empty expression plasmid. After 48 hours, strong *Gm15441* expression was seen, but had no effect on the RNA levels of GFP or UTR-GFP (**Figure 7B**). In contrast, GFP protein translated from the *Txnip* 5’UTR-GFP template was significantly reduced in cells co-transfected with *Gm15441* expression plasmid (**Figure 7C**). This 5’UTR-dependent decrease in GFP protein was most striking when examined by fluorescent microscopy (**Figure 7D and E**). These findings suggest *Gm15441* may regulate TXNIP translation through IRES sites found within its 5’UTR. To examine whether lncRNA *Gm15441* impacts expression of genes that flank *Gm15441* on mouse chromosome 3, namely, *Hfe2 (Hjv)*, *Pol3gl* and *Ankrd34a*, Hepa-1 cells and NIH3T3 mouse embryonic fibroblast cells were transfected with *Gm15441* expression vector, or empty plasmid. The *Gm15441* transgene was expressed and significantly downregulated *Txnip* without impacting the expression of *Hfe2*, *Pol3gl*, or *Ankrd34a* (**Figure 7F**). Together, these data support a regulatory mechanism whereby PPARA induces *Gm15441*, which in turn attenuates inflammation through TXNIP and the NLRP3 inflammasome pathway (**Figure 7G**).

**Figure 7.**
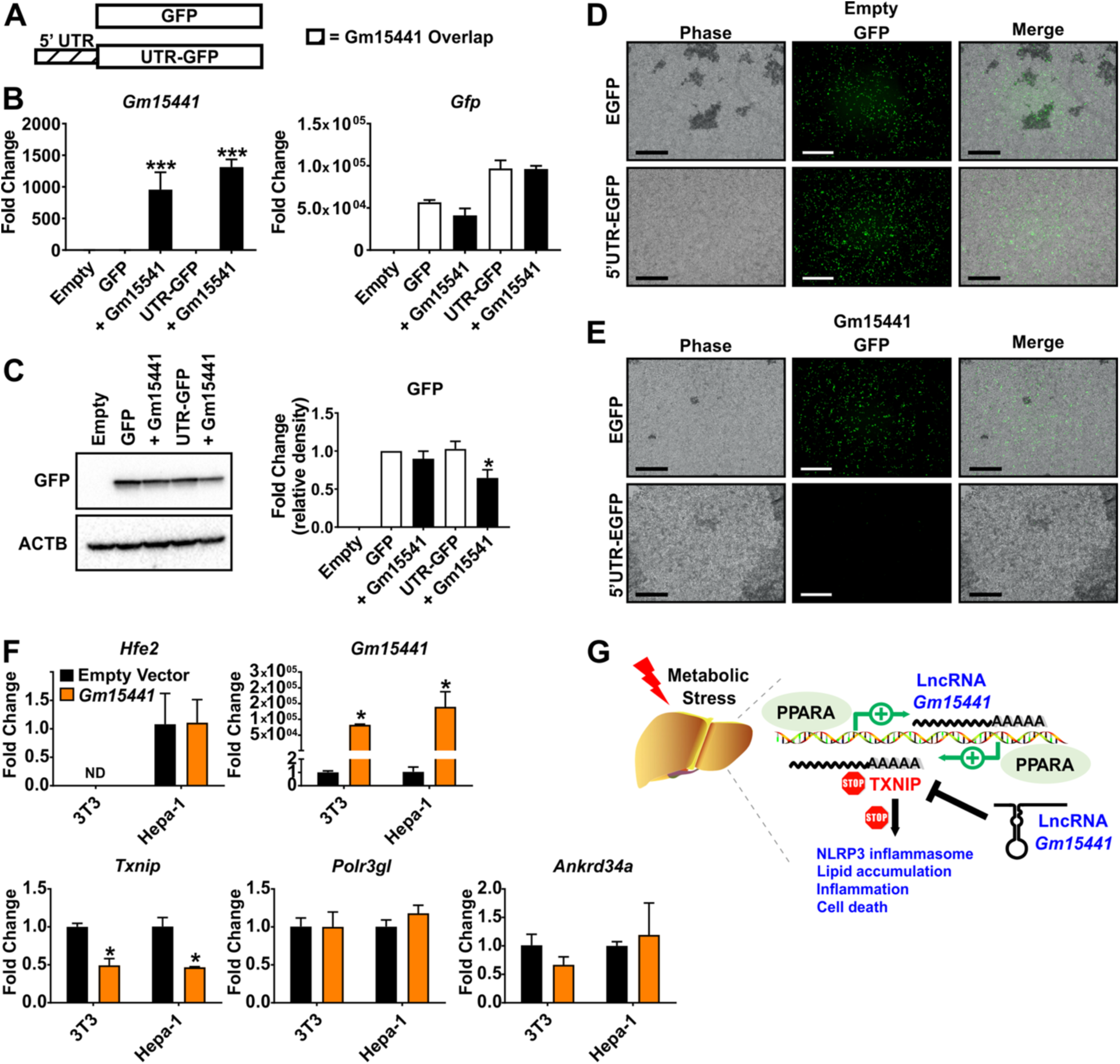
*Gm15441* regulates TXNIP translation in part through IRES sequences found within the 5’ UTR of TXNIP. (A) Schematic construct of inserts for TXNIP of non-5’UTR and 5’UTR sequence containing GFP. (B) Analysis of *Gm15441* and *Gfp* mRNA in Hepa-1 cells. (C) Analysis of GFP protein and relative density of GFP in Hepa-1 cells (*right*). (D and E) Fluorescence of GFP in Hepa-1 cells transfected with EGFP or 5’ UTR sequence containing EGFP with empty or *Gm15441* plasmid DNA for 48 hours. Scale bars represents 20 nm (100x). (F) Analysis of *Hfe2*, *Pol3gl* and *Ankrd34a* mRNAs from Hepa-1 and NIHT3T cells transfected with empty or *Gm15441* expression vector for 24 hours. Each data point represents mean ± SD for n = 3 replicates. *P < 0.05; ***P < 0.001 for comparisons in the absence of Gm15441 (B, C) or to empty vector (F). (G) Model for role of GM15441 in suppressing TXNIP-mediated inflammasome activation.

## Discussion

The physiological alterations that accompany fasting impart several health benefits, including anti-inflammatory effects (Goodpaster and Sparks, 2017; Montagner et al., 2016). As such, the underlying molecular pathways modulated by caloric restriction may present promising new therapeutic targets. PPARA activation during fasting is a key regulatory event of lipid and glucose metabolism. A growing body of evidence indicates that PPARA activation also potently suppresses inflammation in several disease models and tissues (Abcouwer, 2013; Krysiak et al., 2011; Lee et al., 2007; Tomizawa et al., 2011; Zhao et al., 2017). However, the mechanism by which PPARA modulates metabolic stress-induced inflammation is not known. PPARA is activated by endogenous fatty acid metabolite ligands in response to fasting and promotes the uptake, utilization, and catabolism of fatty acids by regulating a wide range of genes that reprogram metabolic pathways to facilitate the use of lipids as an energy source. The direct regulation of protein-coding genes by PPARA is well characterized, but it was not known whether lncRNAs – which may influence gene expression through a variety of mechanisms (Li et al., 2017b; Militello et al., 2018; Zhao et al., 2016) – also serve as direct targets contributing to physiological changes induced by PPARA activation. This possibility was suggested by the finding that several hundred liver-expressed lncRNAs are dysregulated in livers exposed to xenobiotic agonist ligands of the nuclear receptors CAR and PXR (Lodato et al., 2017), which contribute to the regulation of lipid metabolism and whose gene targets overlap with those of PPARA (Cui and Klaassen, 2016; Wada et al., 2009). The present study identified several hundred PPARA-responsive liver-expressed lncRNA genes, including anti-sense lncRNAs, which often contribute to the regulation of genes on the opposing strand (Guil and Esteller, 2012; Qu et al., 2019; Zhang et al., 2017a). Further, a novel PPARA-dependent regulatory axis involving one such anti-sense lncRNA, *Gm15441*, was characterized. *Gm15441* was shown to be transcribed in a liver-specific, PPARA-dependent manner to yield an anti-sense lncRNA that protects against metabolic stress by suppressing PPARA induction of the opposing, sense transcript, TXNIP, and thereby suppresses TXNIP-mediated NLRP3 inflammasome activation.

To ascertain whether PPARA regulates lncRNAs, RNA-seq was performed on livers from wild-type and *Ppara*^−/−^ mice treated with the specific PPARA agonist WY-14643. More than 400 liver-expressed lncRNAs, including many novel non-coding RNA transcripts, were significantly upregulated or downregulated in mouse liver 48 hours after WY-14643 treatment. Only six of these lncRNAs were similarly responsive to WY-14643 in *Ppara*^−/−^ mice, thus establishing the striking PPARA dependence of these non-coding RNA transcriptomic responses. PPARA-dependent, liver-specific expression was verified for six non-coding RNAs, and detailed functional studies were carried out on one such gene, *Gm15441*, which showed an unusually strong induction in WY-14643-treated liver. Importantly, the genomic orientation of *Gm15441* is antisense to that of *Txnip*, which codes for a ubiquitously-expressed protein that facilitates cellular responses to oxidative stress and inflammation (Watanabe et al., 2010). TXNIP was originally identified as a negative regulator of thioredoxin (Muoio, 2007), and subsequent studies found that TXNIP contributes to a wide array of processes in several tissues. Notably, TXNIP causes an increase in intracellular reactive oxygen species generation (Hong et al., 2016) that may inhibit hepatocellular carcinoma cell proliferation (Li et al., 2017a). TXNIP is key regulator of NLRP3 inflammasome activation, which plays an important role in liver fibrosis and hepatocellular carcinoma (Ringelhan et al., 2018; Wree et al., 2017), and its activation is associated with the NLRP3 inflammasome pathway in human diseases (Bai et al., 2019; Kim et al., 2019; Li et al., 2019). TXNIP has also emerged as an important glucose sensor that regulates glucose uptake in response to insulin (Waldhart et al., 2017) and plays an important role in metabolic stress (Mandala et al., 2016; Wu et al., 2013). Another study found that PPARA activation by fenofibrate downregulated *Txnip* mRNA and TXNIP protein in endothelial cells (Deng et al., 2017), suggesting the *Gm15441*-*Txnip* regulatory axis described here for liver may also function in extrahepatic tissues. An earlier study found that *Ppara* mRNA and PPARA target gene mRNAs were elevated in *Txnip*-null mice, and that *Txnip* expression attenuated PPRE-luciferase response when co-transfected with PPARA/RXR and treated with WY-14643 (Oka et al., 2009), suggesting a *Txnip-Ppara* feedback regulatory loop that may also involve *Gm15441*.

LncRNAs regulate diverse cellular processes, including metabolism-related genes expressed on the opposing strand (Marin-Bejar et al., 2017; Villegas and Zaphiropoulos, 2015; Zhang et al., 2017a). Antisense lncRNA GLS-AS is dysregulated under nutrient stress which leads to MYC elevation and stabilization (Deng et al., 2019). The present results showed that *Gm15441* negatively regulates *Txnip* expression during metabolic stress. LncRNA *Gm15441* is antisense to *Txnip*, whose mRNA is decreased in liver coincident with the robust increase in *Gm15441* expression beginning 3 hours after activation of PPARA by WY-14643. The underlying PPARA regulatory interactions controlling activation of the *Gm15441* promoter region were investigated, and four of seven PPARA binding sites identified within 10 kb of the *Gm15441* TSS were confirmed by ChIP assays and shown to be functional in reporter gene assays. Thus, the TXNIP anti-sense lncRNA *Gm15441* is a direct target of PPARA in WY-14643-stimulated mouse liver.

Pharmacological activation of PPARA deregulates NRLP3 inflammasome activity in a mouse model of diabetes (Deng et al., 2017) and prolonged hepatic NLRP3 inflammasome activation leads to hepatocyte death, inflammation, and fibrosis. Thus, the TXNIP-mediated NLRP3 inflammasome pathway was examined. *Txnip* mRNA was significantly decreased by PPARA activation in livers of wild-type, *Gm15441*^+/+^ mice, while TXNIP protein was elevated in livers of *Gm15441*^LSL^ mice. Furthermore, TXNIP-mediated NLRP3 inflammasome activation resulted in cleavage of CASP1 to its mature, proteolytically active form. Mature CASP1 subsequently cleaves target proteins including the proinflammatory cytokine IL1B to its corresponding mature, active form. Mature, cleaved CASP1 and IL1B protein were elevated in *Gm15441*^LSL^ mice, indicating an increase in inflammasome activation. These findings indicate that lncRNA Gm15441 suppresses the TXNIP-mediated NLRP3 inflammasome pathway in response to metabolic stress.

PPARA plays an important role in the fasted mouse liver model (Brocker et al., 2018). Hepatic expression of PPARA target genes *Acadm* and *Cyp4a14* was significantly increased by fasting, as was the expression of *Txnip* mRNA, as was seen in livers of *Gm15441*^LSL^ mice as well as *Gm15441*^+/+^ mice compared with fed mice. TXNIP protein was more highly increased in fasting *Gm15441*^LSL^ mice compared to fasted *Gm15441*^+/+^ mice. Fasting *Txnip* mRNA levels are closely associated with lipid and glucose regulation (Szpigel et al., 2018; Waldhart et al., 2017). In addition, TXNIP-mediated NLRP3 inflammasome pathway target protein expression was significantly elevated in livers of fasting *Gm15441*^LSL^ mice compared to *Gm15441*^+/+^ mice. These results support the proposal that pharmacological and physiological upregulation of lncRNA *Gm15441* prevents metabolic stress by suppressing the TXNIP-mediated NLRP3 inflammasome pathway.

PPARA, a key regulator of global lipid homeostasis, modulates fasting-induced lipid accumulation and hepatosteatosis in mice (Brocker et al., 2018). Lipid droplets appeared in livers of fasted *Gm15441*^+/+^ mice but were more widely distributed in livers of fasting *Gm15441*^LSL^ mice compared to fasting *Gm15441*^+/+^ mice. Histological lipid staining revealed considerably larger amounts of lipid accumulation in livers of fasted *Gm15441*^LSL^ mice than in fasted *Gm15441*^+/+^ livers. Serum and liver TG levels and liver CHOL levels were significantly elevated by fasting in *Gm15441*^LSL^ mouse livers. RNA-sequencing analysis revealed that liver damage, inflammation, and fatty accumulation biomarkers such as *Apoa4*, *Bhmt*, *Lcn2*, *Orm2*, *Saa1*, and *Saa2* mRNAs showed significantly higher expression under fasting-induced metabolic stress in *Gm15441*^LSL^ compared to *Gm15441*^+/+^ mouse liver. LCN2 is marker for liver damage and inflammation (Asimakopoulou et al., 2016; Moschen et al., 2017). Hepatic *Lcn2* and *Orm2* mRNAs are both upregulated in the liver in response to IL1B (Sai et al., 2014; Zhang et al., 2014). *Saa1* and *Saa2* mRNAs are acute phase response proteins and biomarkers of inflammation and their protein products become major components high density lipoprotein (HDL) regulated by IL1B (Leclerc et al., 2019; Lindhorst et al., 1997; Olteanu et al., 2014). Moreover, SAA1 and SAA2 potentiate NLRP3 inflammasome activation, which would act as a feed forward much further increasing IL1B cleavage/maturation/inflammation (Chiba et al., 2009; Zhou et al., 2016). APOA4 plays a role in hepatic TG secretion and is upregulated in response to lipid accumulation (Qin et al., 2016; Zhang et al., 2017b). Notably, APOA4 is a regulator of fasting lipid that is associated with TG secretion and HDL cholesterol in type 2 diabetes (Delgado-Lista et al., 2010; Qi et al., 2007). APOA4 was stimulated by inflammation through activation of TNFR2 and NF-kB signaling in kidney tubular cells (Lee et al., 2017). BHMT remethylates homocysteine to methionine using betaine and contributes to methionine, homocysteine, S-adenosylmethionine, and glutathione homeostasis (Perez-Miguelsanz et al., 2017). In hepatocytes, BHMT upregulation prevent ER stress response, lipid accumulation, and cell death (Ji et al., 2007; Ji et al., 2008). Additionally, BHMT was identified as blood marker for acute liver injury and tumoral liver (Ma et al., 2014). The increase in mature/cleaved IL1B in *Gm15441*^LSL^ mice may at least partially explain the increased expression of *Lcn2, Orm2, Saa1* and *Saa2* mRNAs by fasting in *Gm15441*^LSL^ mouse liver. Taken together, these genes are all associated with inflammation/oxidative stress, consistent with the PPARA-dependent attenuation by *Gm15441* of TXNIP-associated hepatic inflammation in response to metabolic stress.

An internal ribosome entry site (IRES) is an RNA element that allows for translation initiation (Lampe et al., 2018). Deregulation of IRES-mediated p53 translation promoted the defective p53 response against DNA damage in human cancer cells (Halaby et al., 2015). ER stress, serum deprivation, and hypoxia-induced human acetyl-CoA carboxylase 1 (hACC1) are controlled by IRES sequences (Damiano et al., 2018). Beclin-1 independent autophagy is promoted through IRES-dependent translation of hypoxia-inducible factor 1α (HIF1α) (Wu et al., 2014). Polypyrimidine tract binding proteins (PTBs) were shown to regulate TXNIP expression during nutrition starvation by binding to IRES sequences within the *Txnip* 5’UTR (Lampe et al., 2018). By binding to these IRES sequences, PTB prevents translation initiation and suppresses TXNIP protein expression. As lncRNA *Gm15441* is antisense to TXNIP and its exon 3 almost entirely overlaps the *Txnip* 5’UTR, *Gm15441* may block IRES sequences needed for translation of TXNIP. The ability of Gm15441 to modulate TXNIP protein levels *via Txnip* 5’UTR IRES sequences was demonstrated *in vitro* by co-transfection of a GFP reporter with a *Gm15441* expression vector. A decrease in GFP expression was dependent on both Gm15441 mRNA and the *Txnip* 5’UTR sequence, indicating that *Gm15441* in part regulates TXNIP by binding to its 5’UTR, presumably by masking functional IRES sites.

A strand-specific *Gm15441* knockout mouse was developed to assess lncRNA function without impacting regulation of its antisense protein-coding partner *Txnip*. In contrast, previous knockout mouse models developed to evaluate TXNIP function disrupt both *Txnip* and *Gm15441* and therefore abolish the unique regulatory loop that exists between these two genes. *Txnip*-null mice show a higher incidence of hepatocellular carcinoma, with approximately 40% of male mice developing hepatic tumors (Kwon et al., 2010; Sheth et al., 2006). However, the *Txnip*-null mice lack exons 1-4, and thus *Gm15441* is also ablated (Oka et al., 2006). Conditional *Txnip*-floxed mice were developed that target exon 1, which results in deletion of *Gm15441* exon 3 (Oka et al., 2009). However, in another study, the ubiquitously-expressed protamine-*Cre* mouse was employed to generate a *Txnip* conditional-null mouse line (Yoshioka et al., 2007). Hepatocyte-specific *Txnip*-null mice also used an exon 1 floxed allele which results in the removal of *Gm15441* exon 2 (Chutkow et al., 2008). TXNIP expression *in vitro* inhibited hepatocellular carcinoma cell proliferation and induced apoptosis (Liu et al., 2017). Thus, prolonged *Gm15441*-mediated suppression of TXNIP may contribute to PPARA agonist induced hepatocellular carcinoma (Cho et al., 2019; Gunes et al., 2018).

In summary, the present study focused on the role of lncRNA as a potential metabolic regulator. Pharmacological and physiological activation of PPARA promoted induction of large numbers of hepatic lncRNAs, a subset of which are also targeted by the nuclear receptors CAR and PXR. *Gm15441* was identified as a liver-specific, PPARA-dependent lncRNA positioned antisense to the pro-inflammatory gene *Txnip*. Gm15441 expression was shown to suppress TXNIP protein by a mechanism involving the blocking of IRES sites within its 5’UTR. By suppressing TXNIP translation, Gm15441 inhibits TXNIP-mediated activation of the NLRP3 inflammasome as well as subsequent CASP1 cleavage and IL1B maturation. Taken together, these results indicate that lncRNA *Gm15441* directly prevents metabolic stress-induced inflammation and represents a novel therapeutic target for the treatment of inflammatory disorders.

## Author contributions

C.N.B. and D.K. conducted the experiments and performed data analysis. C.N.B., D.J.W., and F.J.G. designed the experiments and wrote the paper. C.N.B. and D.K. performed all the mouse experiments. T.M., K.K. and D.J.W. analyzed NGS data and identified WY-14643-responsive liver lncRNAs, and D.J.W. revised and edited the manuscript. T.J.V., S.T. and J.A.B. contributed to the analysis of mouse experimental data. C.N.B. and F.J.G. supervised and coordinated the overall study.

## Acknowledgements

We thank Linda G. Byrd and John R. Buckley for technical assistance with mouse studies. This work was funded by the intramural research program at the National Cancer Institute, National Institutes of Health and by NIH grant R01-ES024421 to D.J.W. C.N.B. was supported in part by the Postdoctoral Research Associate Training (PRAT) program through the National Institute of General Medical Sciences, National Institutes of Health. D.K. was supported in part by a grant from the Korea Health Technology R&D Project through the Korea Health Industry Development Institute, funded by the Ministry of Health & Welfare, Republic of Korea (HI17C2082). The content of this paper is solely the responsibility of the authors and does not necessarily represent the official views of the National Institutes of Health or Boston University. The Alb-Cre mice were provided by Derek LeRoith (Mount Sinai School of Medicine).

## CONTACT FOR REAGENT AND RESOURCE SHARING

Further information and requests for resources and reagents should be directed to and will be fulfilled by the lead contact, Frank J. Gonzalez.

## EXPERIMENTAL MODEL AND SUBJECT DETAILS

### Mouse models

Male 6- to 12-week-old mice were used for all studies and all mouse strains were on the C57BL/6J background and maintained on a grain-based control diet (NIH-31). Mice were housed in light and temperature-controlled rooms and were provided with water and pelleted chow ad libitum. For pharmacological studies, the mice were provided a grain-based control diet or matched diet containing 0.1% Wy-14643 for 24 or 48 hours. For monitoring the time dependence of gene responses, Wy-14643 was dissolved in 1% carboxymethyl cellulose (CMC) solution and orally administered (50 mg/kg in 200 µl) for indicated time points. At the end of the treatment period, the mice were killed by CO_2_ asphyxiation and tissues harvested. For physiological studies, food was removed for 24 h starting shortly after the onset of the light cycle and endpoints collected at the same time the following day. Animals were then killed, and tissue samples harvested for further analysis. Blood was collected by venipuncture of the caudal vena cava. All animal experiments were performed in accordance with the Association for Assessment and Accreditation of Laboratory Animal Care international guidelines and approved by the National Cancer Institute Animal Care and Use Committee.

## METHOD DETAILS

### Generation of *Gm15441*-null mice

SAGE Laboratories (Cambridge, UK) provided design and construction services for the CRISPR/Cas gene targeting technologies used to create a *Gm15441*-null mouse line. The targeting strategy results in the insertion of a floxed cassette containing a transcriptional stop repeat within the first intron of *Gm15441* (NR_040409.1) (**Table S3**). Presence of the cassette prevents *Gm15441* expression. Crossing with a *Cre* mouse line removes the stop cassette and allows *Gm15441* expression to proceed. Microinjection-ready sgRNA, *Cas9* mRNA, and a plasmid donor with a floxed stop cassette were purchased from SAGE Laboratories. The sgRNA, Cas9 mRNA, and plasmid donor were then injected into C57BL/6J mouse embryos by the Transgenic Mouse Model Laboratory at the National Cancer Institute (Fredrick, MD) using the manufacturer’s recommended protocol. Founder animals were genotyped using primer sets in **Table S4**, and all modifications confirmed by targeted sequencing. Homozygous mice were then backcrossed ten times into the C57BL6 background bred out any off-target effects.

### Mouse genotyping

Genomic DNA was extracted from tail using extraction E buffer, TPS and N buffer. 25 µl of E buffer and 7 ul of TPS buffer were added and stored room temperature at 10 min. Then tails were heated at 95°C in 5 min and 25 µl of N buffer was added. Polymerase chain reaction was used for examined of genotype using the following primers: forward: 5’-TGCGAGGCACGATATGGCGA-3’, reverse: 5’-AGCGCACCTGTCACTTTCCTGC-3’. The amplicon with 1200 and 1700 bp were detected in wild-type and *Gm15441*-null mice, respectively.

### RNA isolation, cDNA library construction, and sequencing

RNeasy Plus Mini Kit (Qiagen, Valencia, CA, USA) was used to extract total RNA from livers from four different treatment and control groups: wild-type mice and *Ppara*^−/−^ mice fed either control diet or fed a diet containing 0.1% WY-14643 for 48 hours and euthanized between 1 and 3 PM. RNAs were extracted from n = 9 to 15 independent livers per group, and after quality assessment of RNA by TapeStation 4200 (Agilent, Santa Clara, CA, USA), high quality RNA samples (RIN>9.0) were pooled, as follows, then used to construct stranded RNA-seq libraries from polyA-selected total liver RNA using an Illumina stranded TruSeq mRNA Prep Kit (Illumina, San Diego, CA, USA). Three independent RNA pools were prepared for each of the 4 treatment groups, with each pool comprised of n = 3, 4 or 5 individual liver RNA samples. The libraries were subjected to 126 cycle paired-end sequencing using an Illumina HiSeq 2500 instrument (Illumina) at the NCI-CCR sequencing facility (Frederic, USA) at a depth of 30-42 million read pairs for each of the 12 RNA-seq libraries. For RNA-seq analysis of *Gm15441*^LSL^ mice, total liver RNA was isolated from wild-type mice and *Gm15441*^LSL^ mice fed either control diet or a diet containing 0.1% WY-14643 for 24 hours and killed between 1 and 3 PM. RNAs were extracted from n = 9-12 independent livers per experimental group, analyzed and used to prepare sequencing libraries as described above. Three independent RNA pools were prepared for each of the 4 treatment groups, with each pool comprised of n = 2-3 individual liver RNA samples. Libraries were subjected to 150 cycle paired-end Illumina sequencing at a depth of 13-21 million read pairs for each of the 12 RNA-seq libraries.

### Analysis of sequencing results

Data were analyzed using a custom RNA-seq analysis pipeline (Connerney et al., 2017) as described elsewhere (Lodato et al., 2017). Briefly, sequence reads were mapped to the mouse genome (release mm9) using TopHat2 (v2.1.1) (Kim et al., 2013). Genomic regions that contain exonic sequence in at least one isoform of a gene (exon collapsed regions; (Connerney et al., 2017)) were defined for each RefSeq gene and for each lncRNA gene. HTSeq (0.6.1p1) was then used to obtain read counts for exon collapsed regions of RefSeq genes, and featureCounts (1.4.6-p5) was used to obtain read counts for exon collapsed regions of lncRNA genes. A set of 24,197 annotated mouse RefSeq genes (which includes some RefSeq lncRNAs) and a set of 15,558 liver-expressed lncRNA genes (Lodato et al., 2017; Melia and Waxman, 2019) was considered for differential expression analysis. These lncRNAs include intergenic lncRNAs, as well as lncRNAs that are antisense or intragenic with respect to RefSeq genes, and were discovered using a computational pipeline for lncRNA discovery described elsewhere (Melia et al., 2016) based on 186 mouse liver RNA-seq datasets representing 30 different biological conditions. RefSeq and lncRNA genes that showed significant differential expression following exposure to WY-14643 were identified by EdgeR as outlined elsewhere (Melia et al., 2016). Genes dysregulated with an expression fold-change (i.e., either up regulation or down regulation) >2 at a false discovery rate (FDR), i.e., an adjusted P-value <0.05 were considered significant and are shown in Table S1 and Table S2. Raw and processed RNA-seq data are available at GEO (https://www.ncbi.nlm.nih.gov/gds) accession numbers GSE132385 and GSE132386.

### Cell culture

Primary hepatocytes were isolated from C57BL6N mice as previously reported (Seglen, 1976) and seeded on collagen-coated 12-well plates (Becton Dickinson and Company, Franklin Lakes, NJ) at a density of 2 x 10^5^ cells in Williams’ Medium E (Thermo-Fisher Scientific, Waltham, MA) supplemented with 5% FBS and penicillin/streptomycin/amphotericin B solution (Gemini Bio-products, West Sacramento, CA). Hepa-1 mouse hepatoma cells and NIHT3T mouse embryonic fibroblast cells (ATCC, Manassas, VA) were maintained at 37°C in a humidified atmosphere of 5% CO_2_ in Dulbecco’s Modified Eagle Medium (DMEM) containing 10% Fetal Bovine Serum (FBS) and 1% of penicillin/streptomycin mixture (Invitrogen, Waltham, MA).

### Histological staining

Fresh liver tissue was immediately fixed in 10% phosphate-buffered formalin for 24 h and then processed in paraffin blocks. Four-micrometer sections were used for H&E staining. Sections were processed by HistoServ, Inc. (Germantown, MD). Slide imaging was performed using a Keyence BZ-X700 series all-in-one microscope with both 20× and 40× objectives, 200× and 400× magnification, respectively.

### Serum biochemistry

Blood was collected from mice and transferred to BD Microtainer Serum Separator Tubes (Becton Dickinson, Franklin Lakes, NJ). Serum was flash frozen in liquid nitrogen and stored at −80C. Serum chemistry analysis for total cholesterol (CHOL), TG was performed using Wako Clinical Diagnostics kits (WakoUSA, Richmond, VA). Serum alanine aminotransferase (ALT) and aspartate aminotransferase (AST) levels were measured using Catachem VETSPEC Kits as recommended by the manufacturer (Catachem, Oxford, CT). Blood GLU levels were measured using a Contour blood GLU meter (Bayer, Mishawaka, IN).

### Western blot analysis

Whole cell extracts were prepared from mouse liver tissue or mouse primary hepatocytes using RIPA buffer supplemented with Halt Protease and Phosphatase Inhibitor Cocktail (Thermo-Fisher Scientific) and 1 mM PMSF. Protein concentrations were determined using the Pierce BCA Protein Assay Kit (Pierce, Rockford, IL). Forty µg of protein was loaded per lane on a 4-12% Criterion TGX Precast Gel (Bio-Rad) then transferred to PVDF membranes using a Trans-Blot Turbo Transfer System (Bio-Rad). Membranes were blocked in 5% nonfat milk followed by an overnight incubation with primary antibodies targeting TXNIP (Novus; NBP1-54578), CASP1 (Proteintech; 22915-1-AP), IL1B (Cell Signaling; #12242), or TUBA1B (Epitomics; 1878-1) at 4°C. Following primary antibody incubation, the blots were washed and incubated with HRP-conjugated secondary antibodies for one hour (Cell Signaling; #7074S, #7076S). The blots were then stripped using Restore Western Blot Stripping Buffer (Thermo-Fisher Scientific) and re-probed with alternate antibodies. An antibody against ACTB (Cell Signaling; 8457L) was used as a loading control. Blot imaging was performed on a ChemiDoc MP System (Bio-Rad) after exposing the blot to Clarity Western ECL Blotting Substrate (Bio-Rad). Protein expression was quantitatively analyzed using band density and ImageJ software (NIH, Bethesda MD) (Schneider et al., 2012).

### Quantitative reverse transcription PCR assays

Total RNA was isolated from fresh mouse liver, mouse primary hepatocytes, and Hepa-1 cells using TRIzol Reagent (Thermo-Fisher Scientific, Waltham, MA, USA) and quantified using a NanoDrop Spectrophotometer (NanoDrop Products, Wilmington, DE, USA). Total RNA (2 µg) was reverse transcribed using All-in-One cDNA Synthesis SuperMix (BioTool, Houston, TX, USA). qRT-PCR analysis was performed using SYBR Green qPCR Master Mix (BioTool). Primers were designed for gene specificity and to cross exon-exon junctions using Primer-BLAST (www.ncbi.nlm.nih.gov/tools/primer-blast/) and purchased from IDT DNA Technologies (Coralville, IA, USA) (**Table S4**). qRT-PCR experiments were designed and performed according to Minimum Information for Publication of Quantitative Real-Time PCR Experiments (MIQE) guidelines (Plain et al., 2014). Results are normalized to actin expression. Values given are fold over control or relative expression value, where appropriate, calculated using the 2ΔCt QPCR calculation method (Pfaffl, 2001).

### Luciferase reporter assays

For luciferase assays, pSG5-PPARA (mouse) and pSG5-RXRA (mouse) were used for transcription factor expression (Shah et al., 2007). Custom GeneBlocks (IDT DNA) were synthesized containing the predicted PPRE sites for Gm15441. GeneBlocks were digested and purified using a Qiagen PCR Purification Kit (Qiagen, Valencia, CA), and cloned into the pGL4.11 for PPRE reporter constructs (Promega, Madison, WI) using a BioRad Quick Ligation Kit (BioRad, Hercules, CA, USA). Prior to performing assays, all constructs were confirmed by Sanger sequencing at the NCI Center for Cancer Research Genomics Core. The phRL-TK renilla luciferase construct was used as a control to normalize for transfection efficiency. Primary hepatocytes were seeded into 12-well plates (4 × 10^4^ cells/well). PPRE reporter constructs were co-transfected into hepatocytes with PPARA and RXR expression. vectors. PPRE-luc plasmid containing an Acox1 PPRE site repeat was used as a positive control (Kliewer et al., 1992). Empty pGL4.11 plasmid was used as negative controls. Plasmids were transfected using Lipofectamine 3000 Reagent (Thermo-Fisher Scientific). Luciferase activities were measured and plotted relative to lysate protein concentrations using the Promega Dual Luciferase Reporter (Promega) assays according to the manufacturer’s protocol. Measurements were taken on a Veritas microplate luminometer (Turner Biosystems, Sunnyvale, CA, USA).

### Chromatin immunoprecipitation

Chromatin was prepared from hepatocytes for ChIP assays as previously described (Kim et al., 2014). Cells were fixed with 4% paraformaldehyde for 15 min, then glycine was added to a final concentration of 0.125 M and incubated for 10 min before harvesting. Chromatin was sonicated using a Bioruptor Pico (Diagenode, Denville, NJ, USA). Chromatin preparations were subjected to ChIP using a ChIP-IT High Sensitivity Kit and Protein G Agarose Prepacked Columns (Active Motif, Carlsbad, CA, USA) using either PPARA (Abcam; Ab24509) antibody. Normal rabbit IgG (Cell Signaling Technologies; #2729S) and Histone H3 (Cell Signaling Technologies; #4620) antibody were used as negative and positive controls, respectively. DNA was purified and concentrated using MinElute Reaction Cleanup columns (Qiagen). qRT-PCR and conventional PCR were performed using 2 µl of ChIP DNA samples from the 50 µl of purified samples using gene-specific primers (**Table S4**). Cycle threshold (Ct) values of ChIP and input samples were calculated and presented as fold change.

### RNA fluorescent in situ (FISH) hybridization

For fluorescence in situ hybridization (FISH) staining of *Gm15441*, *Gm15441*^+/+^ and *Gm15441*^LSL^ mice were fed either a control diet or diet containing WY-14643 for 36 hours. Livers were harvested, fixed in 4% of paraformaldehyde overnight then sent to the Molecular Pathology Laboratory (PHL) at the National Cancer Institute for processing. RNA FISH experiments were performed by the using custom RNAscope probes and reagents developed by Advanced Cell Diagnostics (Newark, CA). Proprietary FISH probes targeted a region of *Gm15441* that does not overlap with *Txnip* to prevent signal interference (**Table S5**). Slide imaging was performed using Aperio ImageScope software (Leica Biosystems, Buffalo Grove, IL, USA).

## QUANTIFICATION AND STATISTICAL ANALYSIS

PPARA ChIP-seq data was downloaded from NCBI Gene Expression Omnibus (GSE61817). CHIP-seq data was uploaded to the Galaxy public server at usegalaxy.org to analyze the data (Afgan et al., 2018). More specifically, data were converted to bigwig file format using Galaxy tools. Bigwig files were then visualized using Integrated Genome Browser (version 9.0.0) (Freese et al., 2016). All results are expressed as means ± SD. Significance was determined by t-test or one-way ANOVA with Bonferroni correction using Prism 7.0 software (GraphPad Software, La Jolla, CA, USA). A P value less than 0.05 was considered significant and statistical significance is indicated in the figure legends.

## Supplemental Legends

**Table S1A. The list of differential gene expression analysis of RefSeq genes. (Related to Figure 1)**

**Tab.le S1B. The list of differential gene expression analysis of lncRNA genes. (Related to Figure 1)**

**Table S1C. LncRNAs responsive to activator of PPARA, CAR and PXR in mouse liver. (Related to Figure 1)** Groups #1, #2: lncRNAs that are consistently induced, or consistently repressed, by all 3 nuclear receptors, as indicated; Groups #3, #4: lncRNAs consistently induced or repressed by PPARA and by CAR, but not by PXR; Groups #5, #6: lncRNAs consistently induced or repressed by PPARA and by PXR, but not by CAR; Groups #7, #8: lncRNAs induced or repressed by PPARA but showing the opposite responses to activators of CAR, PXR, or both receptors.

**Table S1D. Pathway analysis of upregulated genes from the WY-14643 treated mice liver. (Related to Figure 1)**

**Table S1E. Pathway analysis of downregulated genes from the WY-14643 treated mice liver. (Related to Figure 1)**

**Table S2. The list of differential gene expression analysis of RefSeq genes from Gm15441+/+ (WT) and Gm15441LSL (KO) mouse liver. (Related to Figure 6)**

**Table S3. Gm15441 donor plasmid sequence. (Related to Figure 4)**

**Table S4. List of primer sequences used in qPCR and cloning**

**Table S5. Gm15441 target sequence used for fluorescence in situ hybridization probes. (RED: Excluded sequence that is antisense to Txnip) (Related to Figure 4)**

## Highlights

PPARA regulates expression of more than 400 liver-expressed long coding RNAs

Gm15441 is PPARA-dependent lncRNA that is antisense transcript to TXNIP

Gm15441 expression suppresses TXNIP-mediated NLRP3 inflammasome activation

Gm15441 attenuates hepatic inflammation against metabolic stress

## In brief

Fasting is known to elicit anti-inflammatory effects. The molecular mechanisms that thwart inflammation during caloric restriction are not well understood and may represent promising new therapeutic targets. Herein, Brocker, Kim and colleagues identify a novel regulatory loop comprised a long non-coding RNA that suppresses expression of a proinflammatory protein.

## Supplemental Information Titles and Legends

Supplemental Information includes 2 figures and 5 tables and can be found with this article online at XXX

## Supplemental Figures and Tables

**Figure S1.**
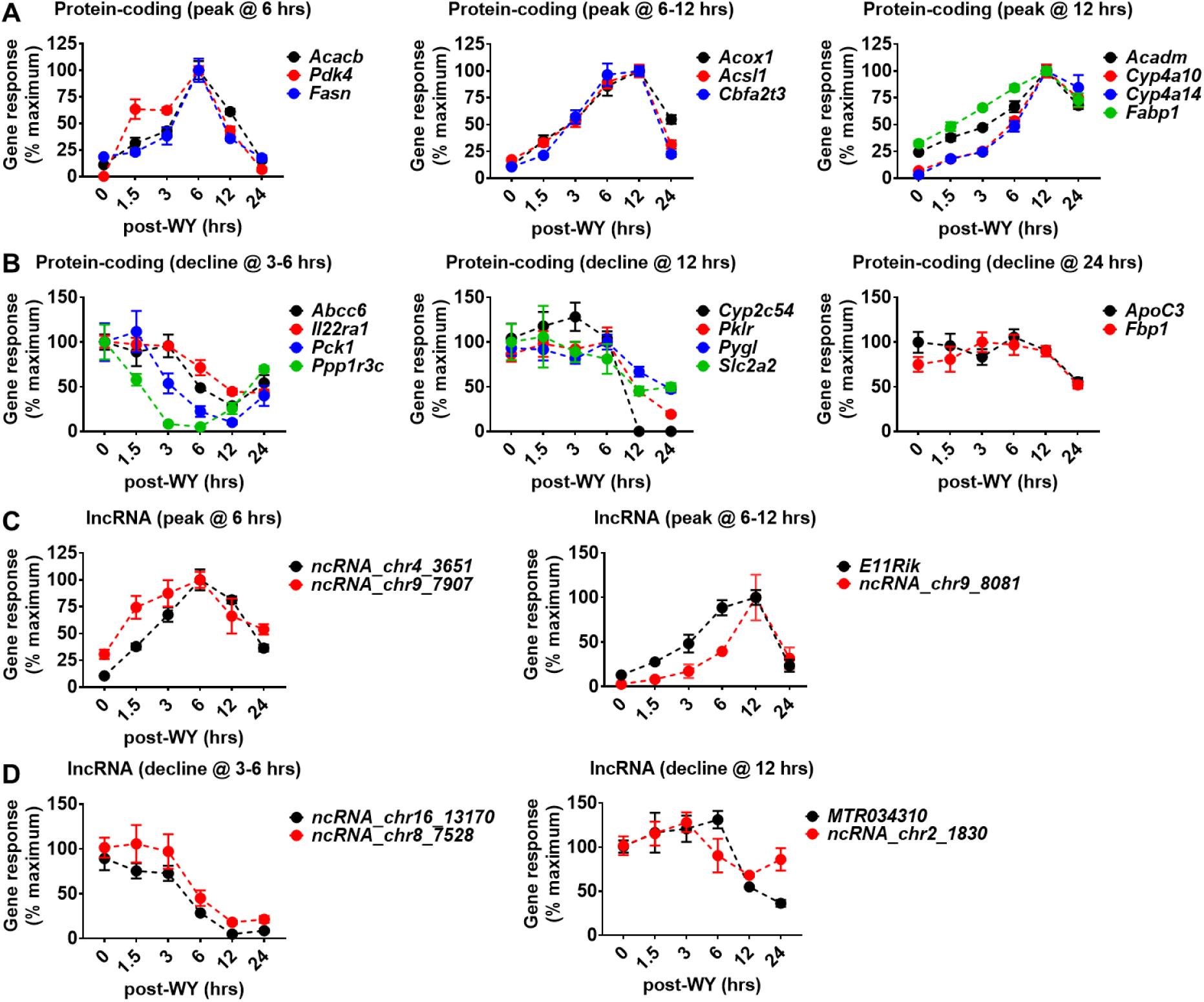
PPARA-dependent lncRNA expression profiles are analogous to protein coding genes. (Related to figure 1) *Ppara*^+/+^ mice were gavaged with the PPARA agonist WY-14643 (50 mg/kg). Livers were collected at t = 0, 1.5, 3, 6, 12, and 24 hours. (A) qRT-PCR analysis of up-regulated protein cording PPARA target genes response by WY-14643 treatment time-dependently. (B) qRT-PCR analysis of down-regulated protein coding gene response by WY-14643 treatment time-dependently. (C) qRT-PCR analysis of up-regulated PPARA target lncRNAs response by WY-14643 treatment time-dependently. (D) qRT-PCR analysis of down-regulated PPARA target lncRNAs response by WY-14643 treatment time-dependently.

**Figure S2.**
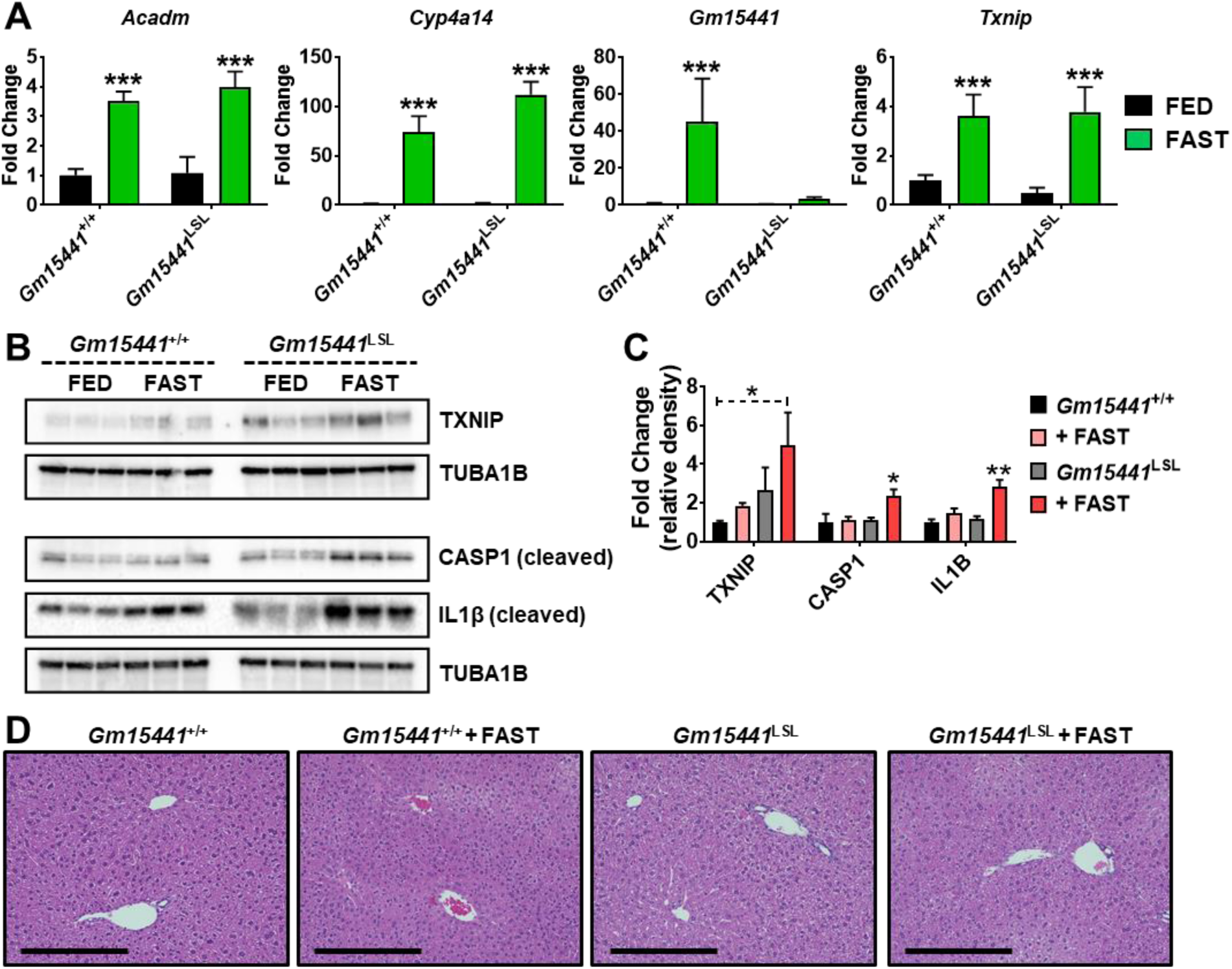
Loss of *Gm15441* potentiates inflammasome activation during physiological response to acute fasting. (Related to figure 6) (A) Analysis of *Acadm*, *Cyp4a14*, and *Txnip* mRNA and *Gm15441* expression from livers of fasted *Gm15441*^+/+^ and *Gm15441*^LSL^ mice, determined by qRT-PCR. (B) Analysis of TXNIP, CASP1 and IL1B protein expression from livers of fasted *Gm15441*^+/+^ and *Gm15441*^LSL^ mice. (C) Densitometric analysis of TXNIP, CASP1 and IL1B protein expression. (D) H&E staining of liver tissues after 24 h fast. Scale bars represents 100 nm (200x). Each data point represents the mean ± SD. *P < 0.05; **P < 0.01; ***P < 0.001.

